# Distinct SARS-CoV-2 populational immune backgrounds induce divergent RBD evolutionary preferences

**DOI:** 10.1101/2023.12.19.572469

**Authors:** Wentai Ma, Haoyi Fu, Fanchong Jian, Yunlong Cao, Mingkun Li

## Abstract

Immune evasion is a pivotal force shaping the evolution of viruses. Nonetheless, the extent to which virus evolution varies among populations with diverse immune backgrounds remains an unsolved mystery. Prior to the widespread SARS-CoV-2 infections in December 2022 and January 2023, the Chinese population possessed a markedly distinct (less potent) immune background due to its low infection rate, compared to countries experiencing multiple infection waves, presenting an unprecedented opportunity to investigate how the virus has evolved under different immune contexts. We compared the mutation spectrum and functional potential of BA.5.2.48, BF.7.14, and BA.5.2.49—variants prevalent in China—with their counterparts in other countries. We found that mutations in the RBD region in these lineages were more widely dispersed and evenly distributed across different epitopes. These mutations led to a higher ACE2 binding affinity and reduced potential for immune evasion compared to their counterparts in other countries. These findings suggest a milder immune pressure and less evident immune imprinting within the Chinese population. Despite the emergence of numerous immune-evading variants in China, none of them exhibited a transmission advantage. Instead, they were replaced by the imported XBB variant with stronger immune evasion since April 2023. Our findings demonstrated that the continuously changing immune background led to varying evolutionary pressures on SARS-CoV-2. Thus, in addition to the viral genome surveillance, immune background surveillance is also imperative for predicting forthcoming mutations and understanding how these variants spread in the population.

## Introduction

According to different strategies for COVID-19 epidemic prevention and control[1–3], the overall infection rate in China was extremely low compared with other countries on December 2022 (e.g., 0.68% in China vs. 49.98% in Israel on Dec 1, 2022, data from www.ourworldindata.org). Meanwhile, China has achieved a relatively high vaccination rate, with 92.54% of the population having received at least one dose of the COVID-19 vaccine and 90.28% having completed vaccination (as of 28 November 2022)[4, 5]. The predominant vaccine used in China was an inactivated vaccine utilizing the original wild-type strain, the elicited antibodies had been largely evaded by the circulating Omicron strains[6, 7]. Moreover, it had been over half a year since the last vaccination for approximately 96% of the vaccinated population. Thus, the Chinese population had a less potent humoral immunity background compared to other countries on December 2022.

In late 2022, China revised its public health control measures[8]. Subsequently, the virus quickly spread across the country and infected over 80% of the population according to an online survey [9, 10]. Given the significant number of infections, there was a growing concern that new variants might emerge within China, akin to how the Delta and Omicron variants originated[11–13]. Although three novel Pango lineages, namely BA.5.2.48, BA.5.2.49, and BF.7.14, were designated based on the genome surveillance data in China[14, 15], a systematic assessment of the mutations, particularly their impacts on immune evasion and ACE2 binding affinity, is missing.

Prior infection and vaccination history gives rise to specific immune response, leading to a phenomenon known as immune imprinting[16], which involves the generation of cross-neutralizing antibodies upon encountering new variants, rather than producing new antibodies[17, 18]. Consequently, the antibody spectrum elicited by the same virus would differ among populations with varying immune backgrounds. This divergence would lead to distinct immune pressure on the virus, which in turn generates variants with different escape mutations. This hypothesis has been indirectly validated through the analysis of differences in the mutation spectrum (the independent occurrences distribution of different mutations) among different SARS-CoV-2 variants circulated at different time periods[19], yet it has not been validated in any particular variant that extensively spread across populations with distinct backgrounds. China’s distinctive immune landscape, combined with the prolonged transmission of the same viral strains in both China and other countries, presents an unparalleled chance to directly scrutinize the evolutionary differences of this virus within distinct immune contexts.

## Results

### Circulation of three SARS-CoV-2 clades in China from December 2022 to March 2023

Between 1st July 2022 and 31st May 2023, a total of 21,346 complete SARS-CoV-2 genome sequences were collected in China after deduplication from GISAID and the RCoV19 databases[20, 21]. We placed all sequences onto the global phylogenetic tree using UShER, following the removal of duplicated sequences and the masking of error-prone positions[22]. We detected three clades predominantly composed of Chinese sequences (constituting over 92% of sequences in the clade). Each clade encompassed more than 1,000 Chinese sequences, and collectively constituting 68.8% of the all sequences from China (Figure 1A). These three clades corresponded to the BA.5.2.48*, BF.7.14*, and BA.5.2.49 lineages, respectively. The most recent common ancestor (MRCA) of the three clades can be traced back to a node that belongs to the BA.5.2 lineage (Figure 1B). The estimated emergence time of the MRCA for the three clades falls within the range of June to August 2022. Hence, the presence of three clades may signify three distinct introduction events, occurring several months prior to the easing of containment measures (Supplementary Fig. S1).

**Figure 1.**
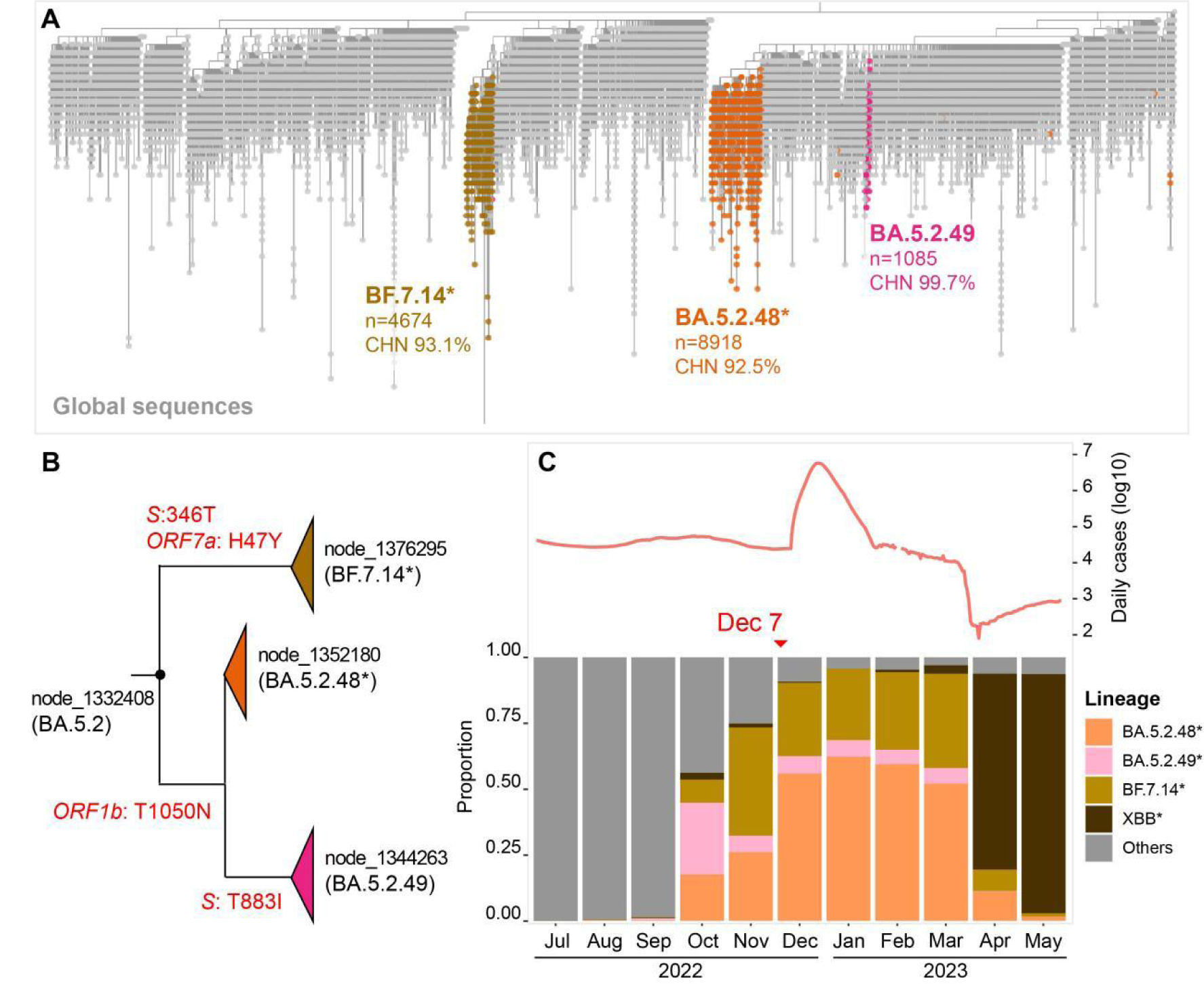
The three SRAS-CoV-2 clades circulating in China. A) The UShER phylogenetic subtree under node 1332408, which is the most recent common ancestor of clades BA.5.2.48, BF.7.14, and BA.5.2.49. Sequences collected in China are colored based on their lineages, and global sequences (collected outside China) are shown in grey. The total number of sequences and the proportion of Chinese sequences in the clade were indicated beneath each clade. B) The phylogenetic relationships between three Chinese-dominant clades. The feature amino acid mutations are labeled on the branch. C) The composition of the circulating SARS-CoV-2 variants in China. The number of daily cases is marked on the top of the panel.

Notably, the three clades had a limited presence in China before October 2022, whereas other clades, including sub-lineages of BA.2, BA.4, and BA.5 were predominant during that period (Table S1). BA.5.2.48*, BF.7.14*, and BA.5.2.49 became the prevailing circulating variants since October 2022, and were supplanted by XBB* in April 2023 (Figure 1C). The daily count of sequences belonging to the three clades exhibited a strong correlation with the number of daily reported cases (Supplementary Fig. S1), and these three clades constituted 93.4% of the sequences during the surge between Dec 2022 and March 2023. Therefore, we opted to investigate the evolutionary dynamics of the SARS-CoV-2 virus using these three clades in subsequent analyses.

### The mutation spectrum in the RBD region differed between China and other countries

We identified 10,692 nucleotide mutation events occurred in the BA.5.2.48* lineage, 7,264 in the BF.7.14* lineage, and 1,219 in the BA.5.2.49 lineage. Considering the small number of sequences and mutation events within the BA.5.2.49 lineage, its sub-lineage association with the BA.5.2.48* lineage (Figure 1B), and the shared receptor binding domain (RBD) sequences with BA.5.2.48*, we consolidated the BA.5.2.48* and BA.5.2.49 lineages for subsequent analyses (named BA.5.2.48/49*).

The distribution of the non-synonymous (NS) mutations in most genomic regions was similar between three Chinese-dominant lineages and their counterparts from other countries (Fig. 2A). And there was a positive correlation between the incidence of NS mutation on BA.5.2.48/49* and those on its immediate predecessor, BA.5.2, in other countries (Supplementary Fig. S2A). Similar tendency was observed between BF.7.14* and its immediate predecessor BF.7, with the exception of the *ORF6* region. Meanwhile, we observed a notable decrease in NS mutations within the *ORF1ab* gene of BA.5.2.48/49* compared to BA.5.2. This reduction was attributed to decreased occurrence of NS mutations in the NSP1, NSP3, and NSP13 regions (Figure 2A, Supplementary Fig. S2B). BF.7.14* exhibited a similar decrease in NS mutations within the NSP13 region when compared to BF.7. Of note, ORF6, NSP1, NSP3, and NSP13 proteins were all involved in innate immune evasion[23, 24].

**Figure 2.**
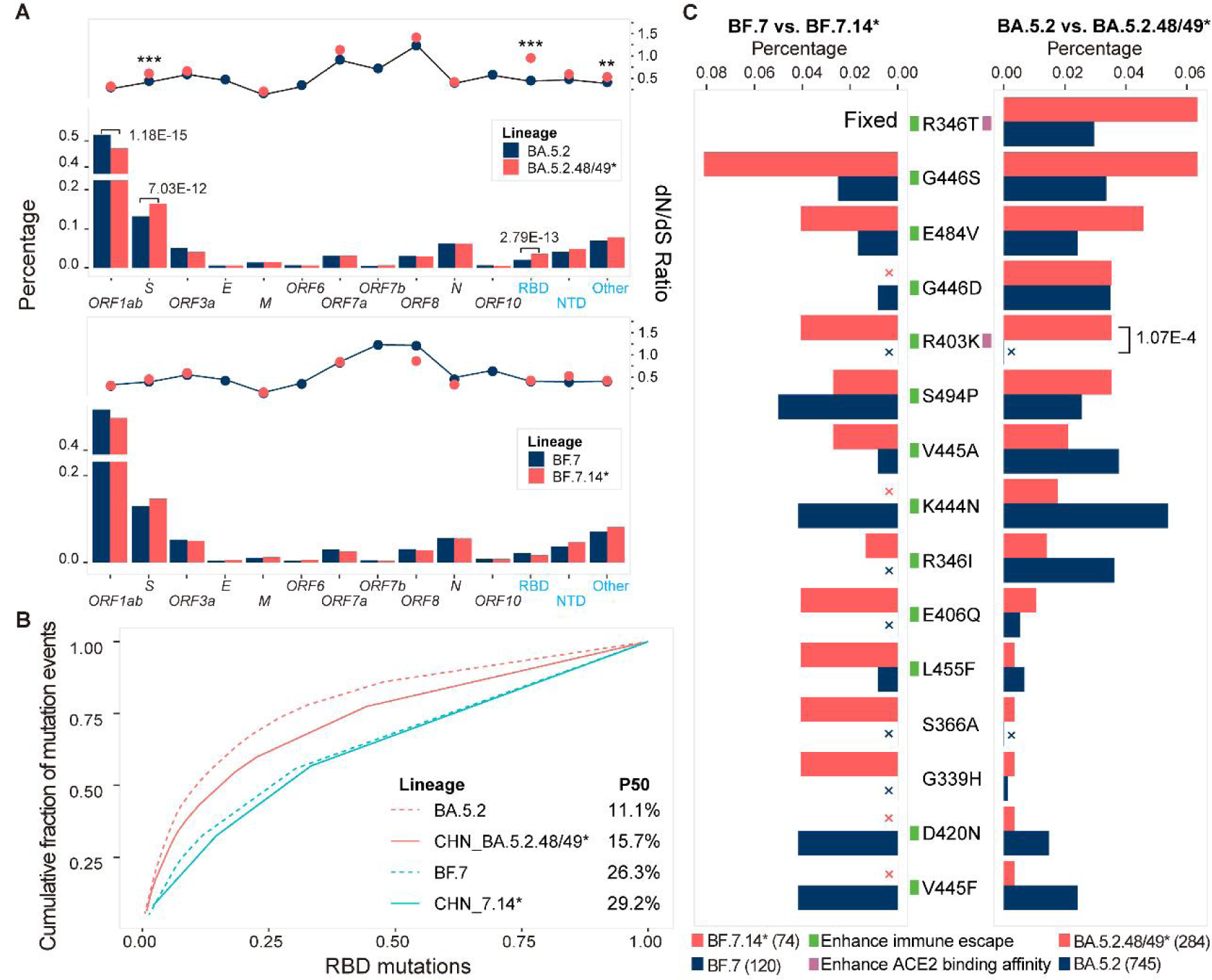
The mutation spectrum differences between variants in China and other countries. A) Mutation distribution across different genes. The sub-lineages of BA.5.2 and BF.7 lineages were not included in the analysis. The bar indicts the proportion of non-synonymous mutations (left y-axis) while the dots indicates the dN/dS ratio for each gene (right y-axis). The ratio for *ORF3a*, *ORF6*, *ORF7b*, and *ORF10* were not shown due to insufficient mutation events number (<100). The Bonferroni adjusted p-value was computed by Fisher’s exact test, with only statistically significant p-values (<0.05) are labeled in the figure. B) The cumulative distribution of RBD amino acid mutations. The x-axis represents RBD mutations sorted by their incidences from high to low. The y-axis represents the cumulative fraction of the mutation events. P50 is the percentage of top prevalent mutations that account for half of the total mutation events. C) The top five most prevalent RBD amino acid mutations in four lineages. Cross indicates no mutation at that position. The green and purple square next to the mutation indicates whether the mutation is able to invade humoral immunity or increase ACE2 binding affinity. The total number of mutation events is provided in the parentheses adjacent to the lineage name underneath the figure. The Bonferroni adjusted p-value was computed by Fisher’s exact test, with only statistically significant p-values (<0.05) are labeled in the figure.

The BA.5.2.48/49* variant also demonstrated an enrichment of non-synonymous (NS) mutations in the S gene, particularly within the receptor-binding domain (RBD) region. This enrichment was associated with a significantly higher dN/dS ratio compared to BA.5.2 (dN/dS: 0.95 *vs.* 0.44, Figure 2A). The distribution of RBD NS mutations in BA.5.2.48/49* was more widely dispersed compared to BA.5.2 (Figure 2B), which may reflect a less concentrated selection pressure on the virus in China. The most prevalent RBD amino acid mutations displayed variations between the BA.5.2.48/49* and BF.7.14* lineages and their international counterparts (Figure 2C). Specifically, BA.5.2.48/49* displayed an enrichment of R346T, G446S, E484V, and R403K mutations compared to BA.5.2. Notably, G446S, E484V, and R403K were also enriched in BF.7.14* when compared to BF.7. Among these mutations, R403K exhibited the most remarkable disparity, and this mutation has been rarely observed in other BA.5 sub-lineages (Table S2). Notably, the R403K is an ACE2 binding-enhancing mutation that ranked 8^th^ in terms of ACE2 binding alterations and 555^th^ in terms of escape scores among all 1,191 possible mutations in the RBD region (Table S3).

### SARS-CoV-2 evolution in China exhibited a preference for heightened ACE2 binding and lower immune evasion

To further elucidate the difference in the driving force behind SARS-CoV-2 evolution in China and other countries, we assessed the impact of RBD amino acid mutations (hereafter referred to as RBD mutations) on two crucial functional aspects—ACE2 binding affinity and immune evasion—that manifested in different countries[19]. We found that RBD mutations occurred on BA.5.2.48/49* had a lower mutation escape score and a higher ACE2 binding score compared to BA.5.2 (Figure 3A). The BF.7.14* showed a similar trend when compared to BF.7.

**Figure 3.**
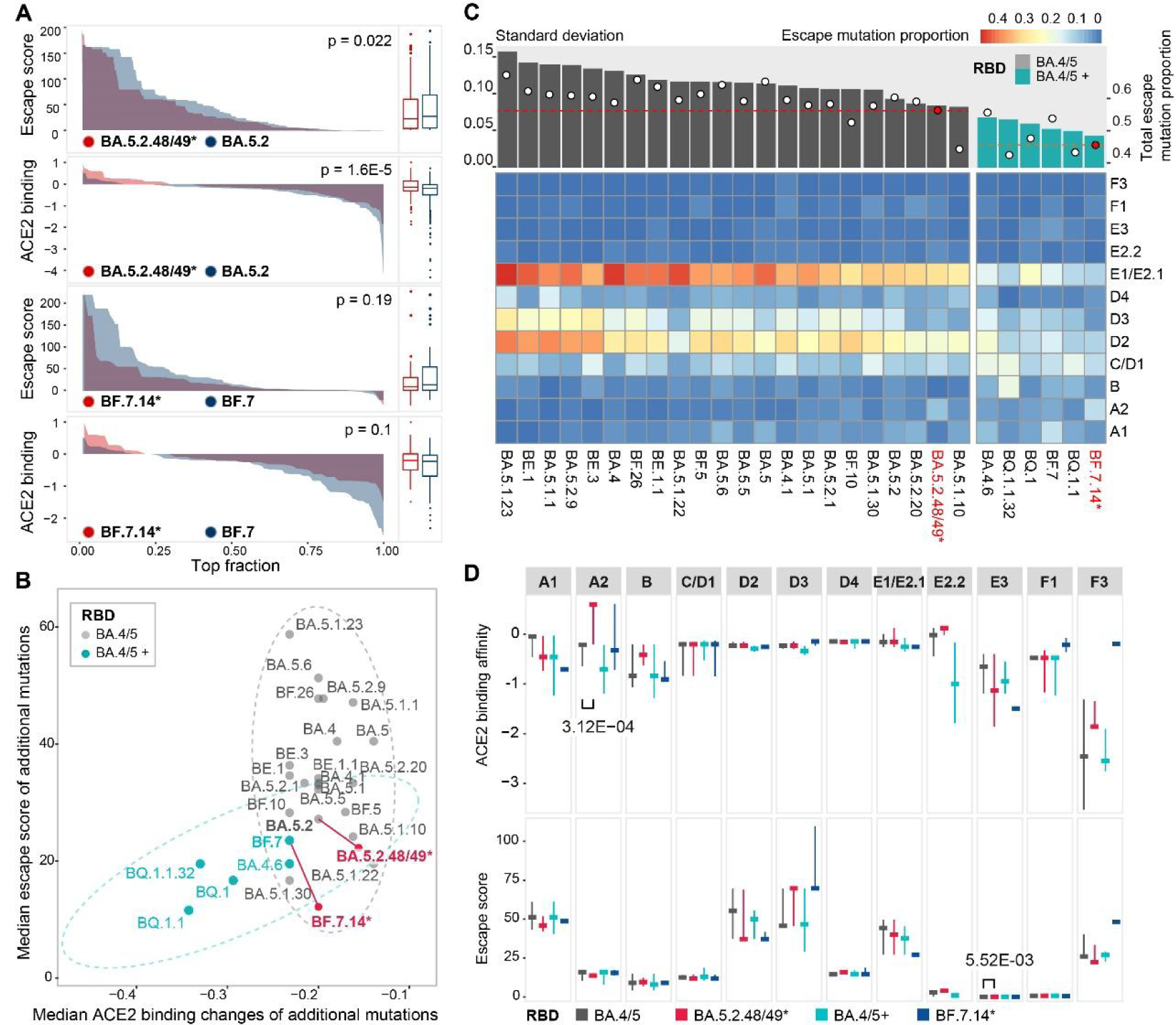
Divergent mutation preferences in viral evolution. A) The comparison of mutation escape scores and ACE2 binding scores between the BA.5.2.48/49* and BF.7.14* lineages and global counterparts (BA.5.2 and BF.7). The x-axis represents the mutations that are sorted by the scores from high to low. The left panel shows the distribution of scores. The right panel shows the box plot of scores. The p-value was calculated by the Wilcoxon rank-sum test. B) The mutation spectrum of BA.4/5 sub-lineages. The mutation functional scores were compared between the BA.5.2.48/49* and BF.7.14* lineages and their close related lineages (BA.4/5 lineages with at least 4600 sequences). The lineages were divided into two groups, the BA.4/5 and the BA.4/5+, based on whether additional amino acid mutations occurred in the RBD region compared to the BA.4/5 prototype. Notably, the BA.5.2.48/49 lineage had no additional RBD mutations whereas the BF.7.14 had one additional mutation R346T. The lineages were positioned based on the median escape score and ACE2 binding affinity score of all mutation events. BA.5.2.48/49* and BF.7.14* lineages are marked in red and connected to their immediate predecessors by a solid line. The circle indicates the 95% confidence interval of two groups. C) The distribution of escape mutations across 12 RBD epitopes. The heatmap illustrates the proportion of escape mutations in each epitope over all mutation events. The percentage of escape mutations in each lineage is denoted by a white circle at the top of the figure, while the red dashed line shows the escape mutation proportion for the BA.5.2.48/49* and BF.7.14* lineages (whose IDs are highlighted in red). The histogram graph depicts the standard deviation of the escape mutation proportion distribution across the 12 epitopes. D) The distribution of ACE2 binding affinity scores and immune evasion scores of escape mutations in 12 epitopes. The horizontal line represents the median value while the vertical line represents the upper and lower quartiles. Comparisons were conducted between BA.4/5 and BA.5.2.48/49*, as well as between BA.4/5+ and BF.7.14*. The Bonferroni adjusted p-values were computed by the Wilcoxon rank-sum test, with only statistically significant p-values (<0.05) are labeled in the figure.

We further extended the analysis by incorporating 24 BA.4/5 sub-lineages that prevalent in other countries with a high number of sequences (>4,600, Table S4). These variants were categorized into two groups (BA.4/5, BA.4/5+) depending on whether additional mutations occurred in the RBD region relative to BA.5 prototype. Interestingly, we found that the two groups can be distinctly differentiated based on their mutation escape scores and ACE2 binding scores (Figure 3B). The group with additional RBD mutations (BA.4/5+) favored mutations with lower immune evasion and lower ACE2 binding potential. This might indicate a reduced selective pressure attributed to the additional mutations in this group. For instance, the R346T mutation in BA.4.6 and BF.7 enhanced the ACE2 binding affinity and facilitated evasion from antibodies targeting the E1/E2.1 epitope; the K444T and N460K mutations within BQ.1 augmented the ACE2 binding affinity and evaded antibodies targeting A1, D2, D3, and E1/E2.1 epitopes[25, 26].

The RBD mutations observed in the BA.5.2.48/49* and BF.7.14* lineages showed a greater propensity for ACE2 binding and a reduced inclination for immune evasion, in comparison to other lineages within the same RBD sequence (Figure 3B). Furthermore, the difference between BA.5.2 and BA.5.2.48/49* was more significant than that between BA.5.2 and other BA.4/5 lineages (Supplementary Fig. S3). Meanwhile, the proportion of immune escaped mutations was considerably lower in the BA.5.2.48/49* and BF.7.14* lineages, and their distribution was more evenly spread across different antigenic epitopes compared to other lineages with the same RBD sequences (Figure 3C). Collectively, these findings suggest a relatively lower and less concentrated immune pressure on the virus in China, while variants acquiring additional binding-enhancing mutations being more prone to spread.

Meanwhile, the distribution of immune escape mutations in the BA.5.2.48/49* and BF.7.14* lineages showed a significant enrichment at the A2 epitope (Supplementary Fig. S4A). However, this did not align with the humoral immune profile acquired from convalescent sera, as BA.5 and BF.7 breakthrough infections induced greater immune pressure on the E1/E2.1 and A1 epitopes, respectively, compared to the A2 epitopes, as opposed to the Omicron reinfection group (mimicking the antibody profile elicited in other countries) (Supplementary Fig. S4B and C). Hence, the proliferation of escape mutations on the A2 epitope was unlikely to originate from immune imprinting or heightened immune pressure on the A2 epitope. Instead, the enrichment in the A2 epitope might be a side effect of enhancing ACE2 binding affinity, as we found that escape mutations in the A2 epitope in BA.5.2.48/49* exhibited an elevated ACE2 binding affinity, while having a minor effect on immune evasion compared to their global counterparts (Figure 3D). Furthermore, we discovered that the A2 epitope was a hotspot for mutations that enhance ACE2 binding affinity, as seven out of the top nine potential ACE2 binding-enhancing mutations were located in this epitope on the BA.5.2 backbone (Supplementary Fig. S4D, Table S3). Among these, four ACE2 binding-enhancing mutations (Q493K, N417I/H, and R403K) were observed in the BA.5.2.48/49* and BF.7.14* lineages, constituting 58% of all mutation events in this epitope region (19/33, Table S5).

### SARS-CoV-2 evolution in China did not generate a potent immune-evading strain

To examine whether immune-evading variants emerged in China, we calculated the remaining neutralization capacity of antibodies identified in convalescent sera from individuals with BF.7* and BA.5.2* breakthrough infections against the newly emerged variants in China. The first immune-evading variant of the BA.5.2.48/49* lineage emerged shortly after its introduction to China in August 2022, while the second immune-evading variant emerged two months later, along with numerous others. The average immune evasion capacity of the circulating variants is limited until January 2023, when highly immune-evading variants emerged, resulting in a 27% reduction in immune pressure (Figure 4A). However, these newly emerged immune-evading variants did not gain significant selective advantage at the population level until May 2023, as the original variant continued to be the predominant variant in the new cases. In contrast, newly emerged variants of BA.5.2 with increased immune evasion capacity exhibited a significant selection advantage since August 2022 in other countries (10 months after the first appearance of BA.5.2). The immune evasion dynamics of the BF.7.14* was similar to that of BA.5.2.48/49* (Figure 4B). Despite the prolonged circulation of immune-evading variants of BF.7* and BF.7.14* circulated in the population (5-10 months), their frequencies in the population did not increase significantly, suggesting no obvious selection advantage over the original variant.

**Figure 4.**
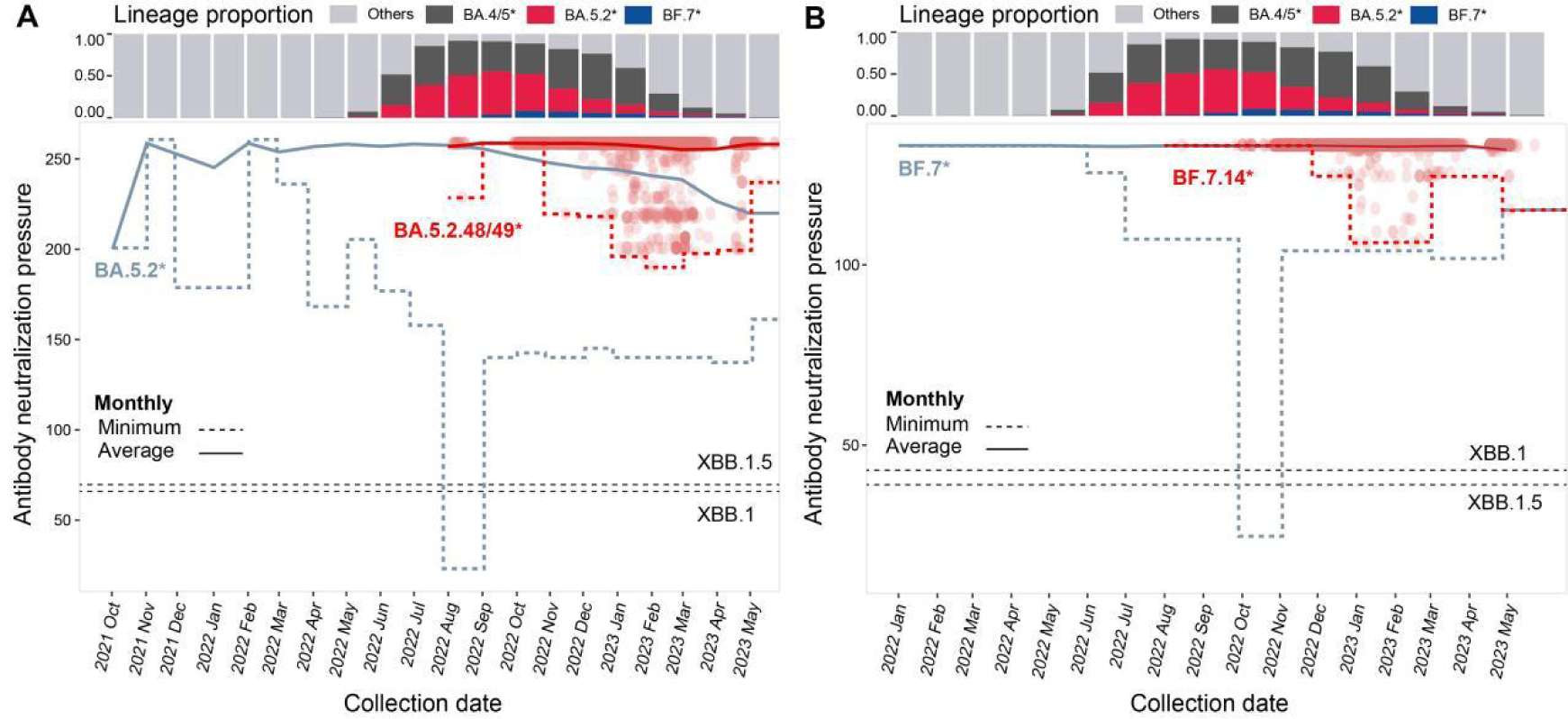
Dynamics of immune evasion capacity during viral evolution. A) Antibody neutralization pressure dynamics on BA.5.2* and BA.5.2.48/49*. B) Antibody neutralization pressure dynamics on BF.7* and BF.7.14*. Each red dot represents a sequence collected in China. Dots representing BA.5.2* and BF.7* sequences are not shown due to the large sample size. The dashed line indicates the minimum values of antibody neutralization pressures across all variants circulating at each time point. The solid line represents the average values of antibody neutralization pressures across all variants circulating at each time points. The global proportion of virus lineages in the infected cases at each time point is displayed on the top of the figure (same for A and B). Notably, BA.4/5* does not include BA.5.2* and BF.7*, and BA.5.2* does not include BF.7*. The antibody neutralization pressures for XBB.1 and XBB.1.5 are depicted as black dashed lines.

The imported XBB lineage replaced the BA.5.2.48/49 and BF.7.14 lineages, emerging as the prevailing variant among newly infected individuals since April 2023. The XBB lineage exhibited a significantly greater immune evasion capacity compared to the newly emerged variants of BA.5.2.48/49, BF.7.14, BA.5.2, and BF.7. Its superior ability to evade the antibodies elicited by prior Omicron variants has also been well documented in recent studies[27, 28].

## Discussion

In this study, we examined the evolutionary trajectory of SARS-CoV-2 in China both during and after the large-scale infection and compared with that of their global counterparts. We found that the BA.5.2.48/49* and BF.7.14* variants, which infected more than one billion individuals, exhibited distinct RBD mutation preferences in contrast to their immediate predecessors, BA.5.2 and BF.7, as well as other Omicron variants sharing the same RBD sequences. The mutations occurring in the RBD region of BA.5.2.48/49* and BF.7.14* variants exhibited three characteristics. 1) The distribution of mutation events was less concentrated; 2) The mutations resulted in a weaker immune evasion capability; 3) The mutations resulted in an elevated ACE2 binding affinity; compared to their global counterparts. Since the variants in China and other countries share the same RBD sequence and nearly identical complete genomes, we speculate that these characteristics were associated with the differences in the immune background between China and other countries.

Due to a low infection rate, a long time since the last vaccine administration, and the mismatch between the vaccine strain and the circulating strain, the humoral immune barrier and immune pressure on the virus at the beginning of the outbreak should be lower in China compared to other countries. This may explain the rapid spread of infections and the reduced occurrence of immune-evading mutations in the BA.5.2.48/49* and BF.7.14* lineages. Meanwhile, because of the less frequent breakthrough infections and reinfections, the virus underwent weaker immune pressure on specific epitope regions compared to other countries that influenced by the immune imprinting effect[29]. This may elucidate why mutations in the BA.5.2.48/49* and BF.7.14* lineages were more widely distributed in the RBD region, and why immune-evading mutations were more evenly distributed across different epitopes.

In a population with a relatively low level of humoral immunity, variants with mutations that enhance transmissibility are more prone to establishing infections, and thus have greater fitness in comparison to variants with immune-evading mutations, which has been observed in our study and previous studies[19, 30, 31]. However, since some mutations, like R403K, could influence both ACE2 binding affinity and immune evasion, the enrichment of ACE2 binding-enhancing mutations in the BA.5.2.48/49* and BF.7.14* lineages had also led to the accumulation of immune-evading capacity within the A2 epitope, which accounts for approximately 11.9% and 8.4% of the estimated immune pressure on BA.5.2.48/49* and BF.7.14*, respectively. This could potentially alter the immune pressure exerted on the virus and result in a divergent mutation trajectory in the future.

It is worth noting that, despite sporadic immune evasion mutations being identified in the viral genome, the immune-evading variants of the BA.5.2.48/49* and BF.7.14* lineages did not exhibit transmission advantage against the original strain in the population until May 2023. This might be attributed to the antibody concentration not having significantly decreased yet, along with the effective cross-protection of antibodies among different variants[32, 33]. For BA.5.2*, it took 10 months that the proportion of immune-evading variants started to increase in the infected population, when the infection proportion of BA.4/5* was approximately 20%; For BF.7*, we have not observed the turning points until May 2023. Thus, evolving a new advantageous variant from an existing strain may require a long time, possibly exceeding one year. However, the emergence of new variants that are not evolutionarily related to previously infected variants, possibly originating from individuals with chronic infections[34], might rapidly replace the previous variants due to their exceptional immune evasion capacities[35], such as the displacement of the Delta by Omicron and the recent replacement of BA.5 by XBB.

Convergent mutations have been frequently observed in various Omicron sub-lineages, and this trend had become more significant over time[19, 29]. However, our results indicate that the intensity and distribution of immune pressure dynamically change along with the emergence of new immune-evading variants, and the trend of convergent evolution became less remarkable in recent lineages (Figure 3). The BQ.1, BA.4.6, and BF.7 lineages exhibited a distinct mutation spectrum characterized by lower immune evasion, reduced ACE2 binding affinity, and a more widely distributed pattern, in contrast to other BA.4/5 sub-lineages that retain the prototype RBD sequence. We speculate that this could be attributed to two factors. On the one hand, the emergence of additional immune-evading mutations had led to a significant reduction in overall immune pressure, particularly in regions targeted by the most potent antibodies. On the other hand, the rising rate of reinfection might undermine the immune imprinting effect and restore some antibody diversity, which is supported by a recent study on individuals re-infected with the Omicron variants[36]. Nevertheless, our understanding of the patterns and trends in the population’s immunity landscape remains limited. It is imperative to establish real-time monitoring and estimation methods for assessing the magnitude and extent of the immune pressure, to accurately predict the future direction of viral evolution.

A limitation of this study arises from the differences between the variants circulating in China and those in other countries. Although they share the same RBD sequences, there are some differences in the *Spike* gene and other non-structural genes (Figure 1B). These dissimilarities could potentially lead to varying immune responses, consequently resulting in distinct immune pressures on the virus. Unfortunately, only a limited number of the three predominant variants in China have been reported in other countries, preventing us from conducting a comparative analysis of these variants in other countries. Nevertheless, the presence of distinct variants circulating across different countries ensures that there has been no transmission of these variants between China and other countries. This in turn guarantees that the mutation and immune backgrounds correspond accurately. Another limitation arises from the limited number of lineages being compared. Including endemic lineages from other countries with relatively strict disease prevention strategies would enhance the reflection of the correlation between immune background and mutation preference. However, we cannot find any other Omicron lineages with more than 4600 sequences that mostly (>90%) collected from any of these countries, highlighting the uniqueness and superiority of our study. Furthermore, while the control lineages/samples used for comparison in this study were collected from different countries, potentially possessing diverse immune backgrounds, all these countries experienced multiple waves of SARS-COV-2 infections during the Omicron era. Consequently, their immune backgrounds were primarily shaped by Omicron variants, rather than being shaped by pre-Omicron variants or prototype vaccines in the Chinese populations, which enables a reasonable and practical comparative analysis in our study.

The immune background inducted by either infection or vaccination is a driving force of virus evolution. Our study has demonstrated a diverse evolution trajectory of SARS-CoV-2 within populations possessing distinct immune backgrounds, shedding light on the emergence and circulation of certain variants in specific geographic regions. In addition to immune pressure, other factors like ACE2 binding affinity, host genetics, and drug usage may also contribute to the evolution of SARS-CoV-2. Quantifying the interplay between these factors and virus evolution to establish a predictive model for the evolution of SARS-CoV-2 remains a substantial challenge we are confronted with.

## Method

### Data Preparation

We retrieved 18,955 complete SARS-COV-2 sequences collected from China from the GISAID database[20] and 10,821 sequences from the RCoV19 database[21], with collection date between 1 July 2022 and 31 May 2023. Sequence ID, sequences, collection date, and submitting laboratory names were used to remove duplicate sequences (8,430), leaving a total of 21,346 sequences for subsequent analyses (Table S1). Daily case numbers in China during the outbreak were retrieved from the OWID website (https://github.com/owid/covid-19-data)[37]. Vaccination information was retrieved from the Our World in Data website (https://ourworldindata.org/coronavirus-data). The estimated global daily number of infections was obtained from a previous study[38], and the accumulative infection rate of a specific variant was calculated by summing the product of daily variant proportion and the estimated daily infection rate.

### Mutation identification and incidence estimation

A deduplication was performed between the 21,346 Chinese sequences and the sequences included in the masked globally SARS-COV-2 mutation-annotated tree (downloaded on 31 May 2023 from http://hgdownload.soe.ucsc.edu/goldenPath/wuhCor1/UShER_SARS-CoV-2/ which contains 7,129,948 public sequences) through metadata comparison. Alignment of the additional Chinese sequences was done by MAFFT[39] (v7.453), and the aligned sequences were placed on the same tree using the UShER script[22, 40], with 481 problematic sites masked[41]. The mutation events were retrieved from the resulting phylogenetic tree using our customized scripts. First, we employed the matUtils tool from the UShER toolkit to transform the protocol buffer format into JSON format. Then, to reduce the number of false positive events caused by incorrect placement of the sequence on the phylogenetic tree, mutation events were exclusively identified within leaf nodes (actual sequences) or internal nodes possessing at least one identical descendant that was a leaf node. Meanwhile, no more than two mutations were allowed between the node and its parental node. Singleton mutations were kept for the analysis. The number of mutation events identified on the phylogenetic tree was used to represent mutation incidence.

When comparing the mutation spectrum between different BA.4/5 lineages, only sub-lineages with more than 4,600 sequences (the number of BF.7.14 sequences) were included for the analysis to minimize the bias caused by a small sample size.

### Searching for SARS-CoV-2 clades in China

We conducted a search for clades primarily composed of sequences from China, stipulating a criterion of having over 80% of sequences originating from China. In total, 1,050 distinct non-overlapping clades were identified, and the three most prominent clades, each comprising over 1,000 Chinese sequences and representing a proportion exceeding 92%, were selected. The BEAST[42] (v2.6.6) was used to infer the TMRCA of each clade. The substitution model was TN93 that selected by the BModelTest function. Visualization of the phylogenetic tree was performed using Taxonium[43].

### Calculation of the ACE2 binding affinity score and the immune escape score

The antibody spectrum, neutralizing activity, antibody epitope group, and raw mutation escape score were obtained from previous studies[29, 36]. Briefly, a total of 1,350 antibodies were identified in the sera of vaccinated individuals and convalescent patients of the wild-type (WT), BA.1, BA.2, BA.5, and BF.7 variants. The impact of mutations in the RBD region on the neutralization effectiveness of antibodies was obtained through a high-throughput Deep Mutational Scanning (DMS) approach. For each mutation, a raw escape score was calculated by fitting an epistasis model that captures the extent of alteration in antibody neutralization effectiveness attributed to the mutation[44]. The raw escape score for each antibody was then normalized to the highest score among all mutations and multiplied by the neutralization value of the antibody. Then, the escape score for a mutation was calculated by summing the escape scores across all antibodies.

The antibodies were classified into 12 epitopes based on their escape profile against the BA.5 variant. For each epitope group, mutations with an escape score (average of the scores against all antibodies belonging to the epitope group) greater than three times the average escape score of all mutations were defined as immune escape mutations.

The ACE2 binding affinity data was obtained from a previous study utilizing a MDS approach[45]. The ACE2 binding affinity score of the mutation was represented as the sum of ACE2 binding value and RBD expression value based on the BA.2 variant.

### Estimation of the antibody neutralization pressure on the variant

The immune pressure exerted on the SARS-CoV-2 variant was calculated by summing the neutralizing activity of all antibodies originating from a specific immune background, i.e., BA.5 convalescents sera for BA.5.2 and BA.5.2.48/49, BF.7 convalescents sera for BF.7 and BF.7.14. When additional mutation emerged within the variant, the updated neutralizing activity of each antibody was calculated employing the formula provided by the SARS-CoV-2 RBD antibody escape calculator[46].

## Data and code availability

All data generated in this study, including original input sequence files and phylogenetic files, as well as all customized scripts were uploaded to the GitHub website along with an introduction (https://github.com/ipplol/SARS2EVO_CHN, DOI: 10.5281/zenodo.8248127).

## Acknowledgments

We gratefully acknowledge all data contributors, i.e., the authors and their originating laboratories responsible for obtaining the specimens, and their submitting laboratories for generating the genetic sequence and metadata and sharing via the GISAID Initiative (EPI_SET ID: EPI_SET_230719ou, doi: 10.55876/gis8.230719ou) and the RCoV19 database, on which this research is based.

## Author contributions

M.L. designed the study. W.M. and M.L. wrote the manuscript with input from all authors. W.M. and H.F. performed bioinformatics analyses. Y.C. and F.J. generated the DMS and neutralization data and supervised the immune analysis.

---

**Supplementary Fig. S1.**
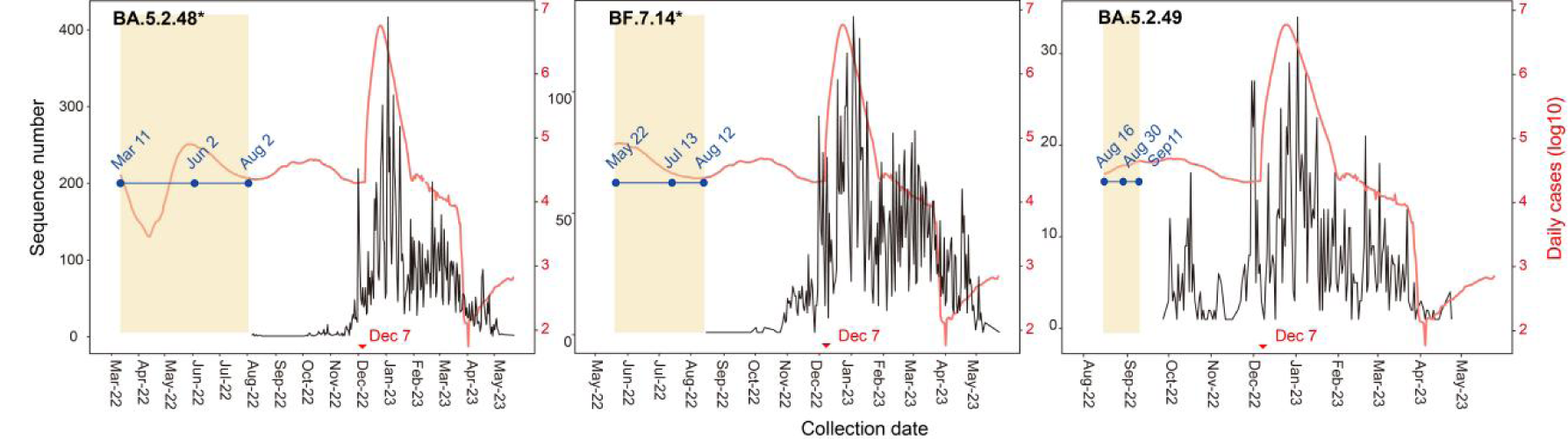
The daily sequence numbers and cases. The red line indicates the daily number of cases in China (right y-axis). The black line indicates the daily number of sequences collected in China that are uploaded to public databases (left y-axis). The occurrence time of the most recent common ancestor for each clade is inferred by BEAST and marked in blue, displaying both the median and the 95% confidence interval. The Pearson correlation coefficient between the number of daily sequence and the number of daily reported cases were 0.48, 0.44, and 0.41 (p<0.0001) for BA.5.2.48*, BF.7.14*, and BA.5.2.49.

**Supplementary Fig. S2.**
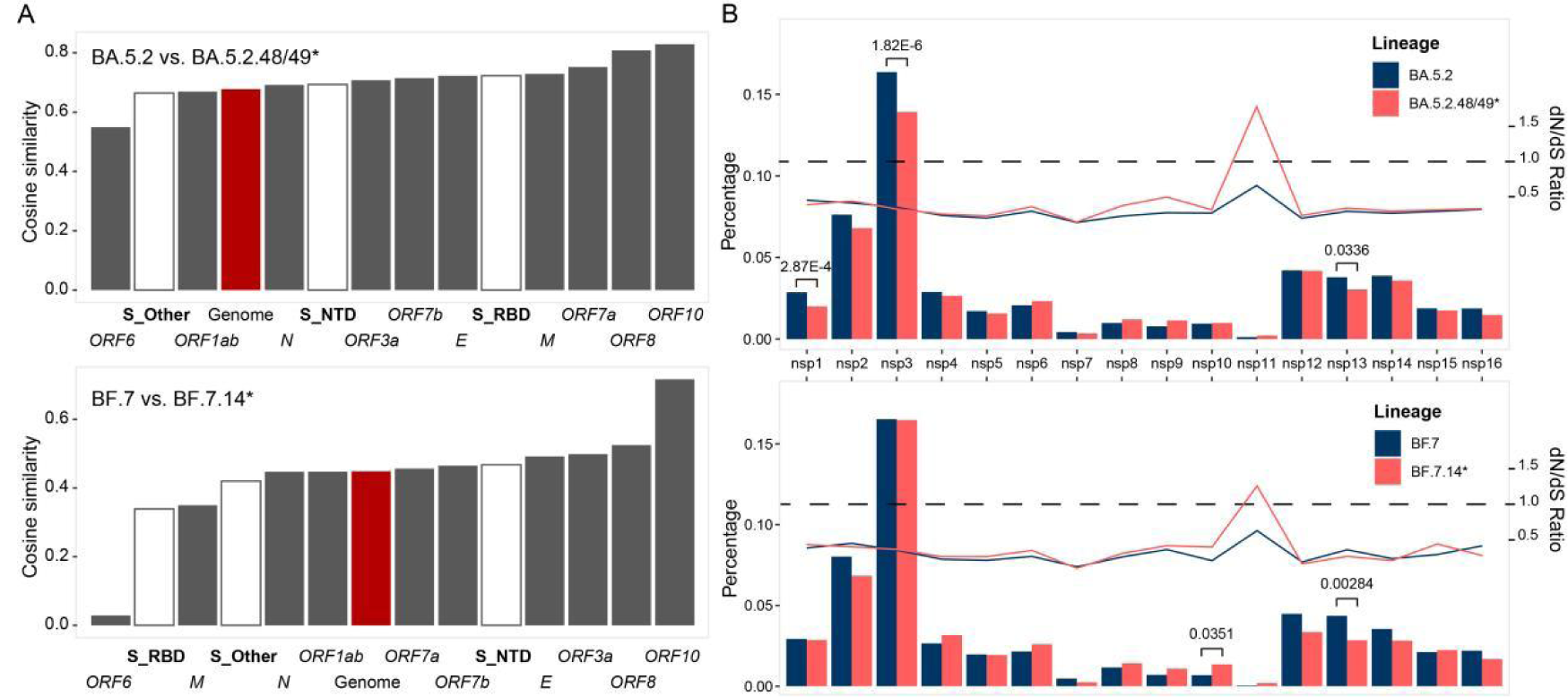
Different in mutation incidence and distribution between the BA.5.2.48/49* and BF.7.14* lineages and their global counterparts. A) Correlation of mutation incidence between the BA.5.2.48/49* and BF.7.14* lineages and their global counterparts. The cosine similarity was calculated based on the incidence of non-synonymous mutations in different genes of SARS-CoV-2. *S* gene was categorized into S_RBD, S_NTD, and S_Other in the analysis. B) Distribution of non-synonymous mutation events across sixteen nonstructural proteins (NSP) regions of the *ORF1ab* gene. The bar indicts the proportion of non-synonymous mutations (left y-axis) while the dots indicates the dN/dS ratio for each gene (right y-axis). The Bonferroni adjusted p-value was calculated by Fisher’s exact test, with only statistically significant p-values (<0.05) are labeled in the figure. The sub-lineages were not included in either the BA.5.2 or the BF.7 lineage.

**Supplementary Fig. S3.**
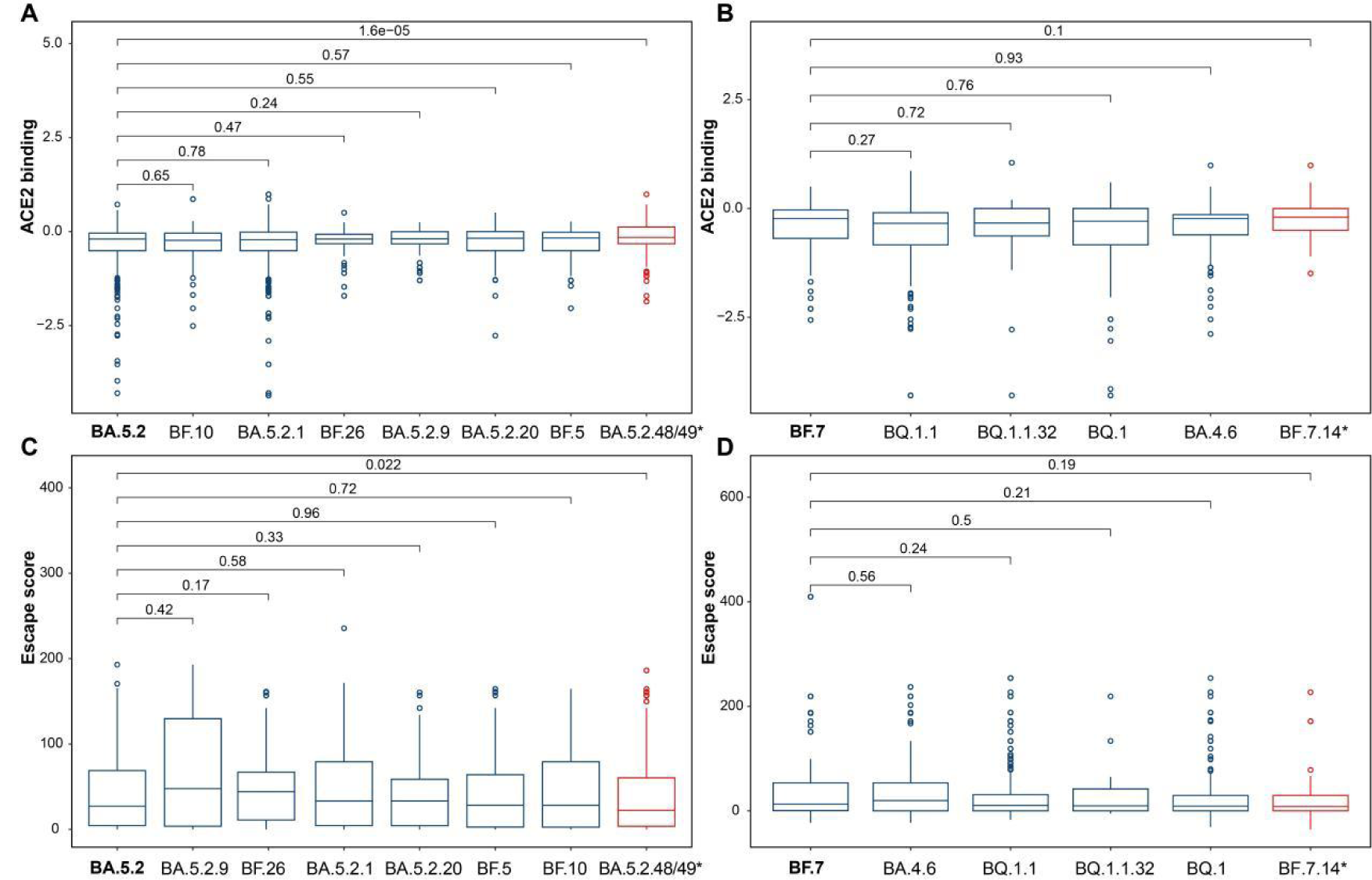
The comparison of mutation escape scores and ACE2 binding scores between BF.7, BA.5.2, and other lineages. A) Comparison of the ACE2 binding scores between the BA.5.2 and other lineages that have the same RBD sequence. B) Comparison of the ACE2 binding scores between BF.7 and other lineages that have additional mutations in the RBD compared to BA.5.2. C) Comparison of the escape score between BA.5.2 and other lineages that have the same RBD sequence. D) Comparison of the escape scores between BF.7 and other lineages that have additional mutations in the RBD compared toBA.5.2. Lineages were sorted by the median value.

**Supplementary Fig. S4.**
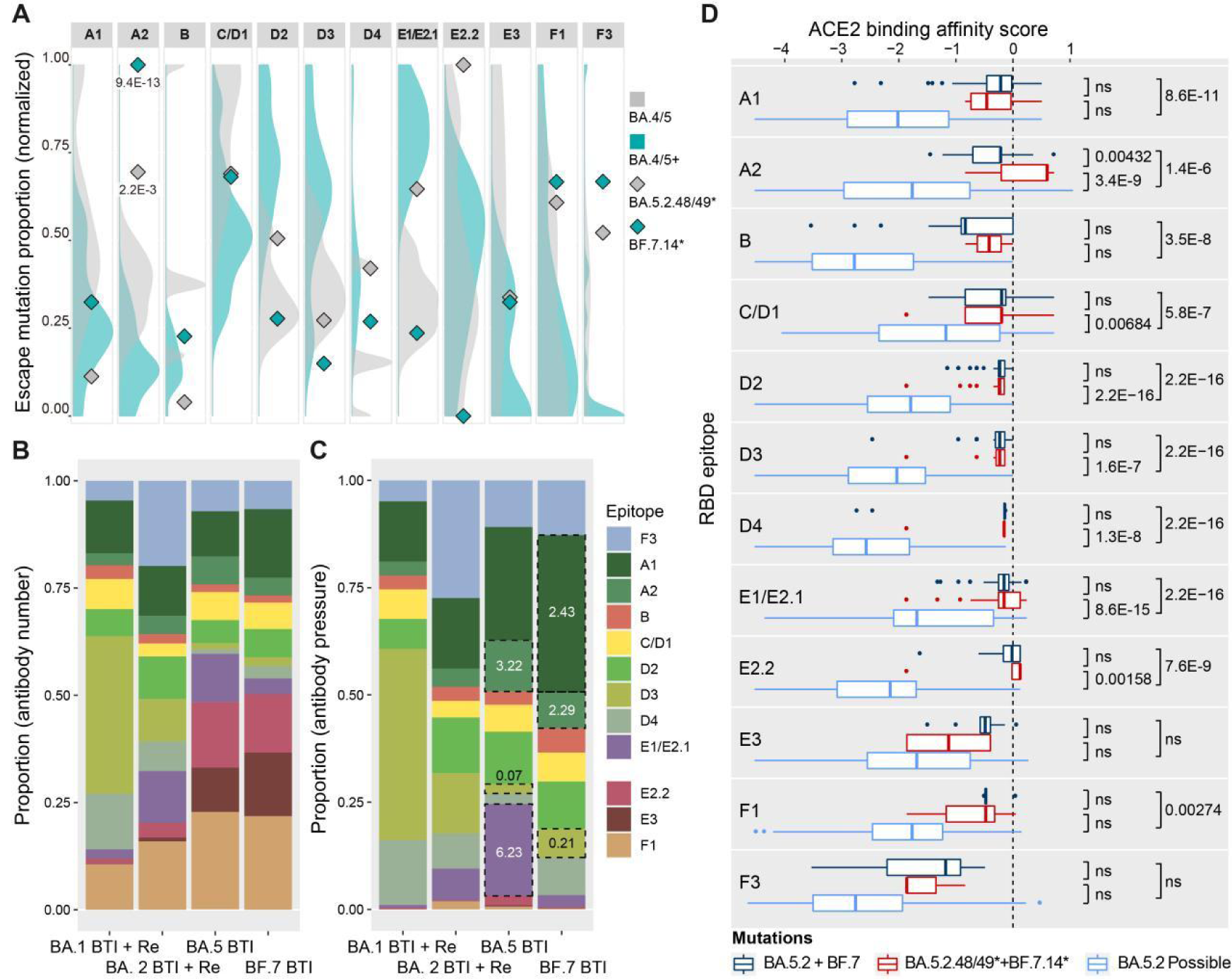
The comparison of the property of escape mutations between the BA.5.2.48/49* and BF.7.14* lineages and global counterparts. A) The proportion of escape mutations in different lineages. The proportion of escape mutations was normalized using the maximum value within each epitope. The density distribution was estimated using data from the BA.4/5 and BA.4/5+ groups, with the exclusion of the BA.5.2.48/49 and BF.7.14 lineages. The significance of the deviation in the escape score of BA.5.2.48/49 and BF.7.14 from the background distribution (assuming a normal distribution) was calculated as the probability of obtaining a value equal to or greater than the observed value, with only statistically significant p-values (<0.05) are labeled in the figure. B) The composition of the antibodies targeting different epitopes in convalescent sera with different infection histories. C) The distribution of humoral immune pressure on different epitopes. Dotted boxes highlight epitopes with immune pressure alterations of over two-fold between convalescent sera from reinfection and breakthrough infection cases (the value within the box denotes the ratio of immune pressure in breakthrough infection sera to that in reinfection sera). Immune pressure on a specific epitope was calculated by summing the normalized neutralization IC50 values of the antibody that target the epitope. BA.1 BTI+Re: BA.1 breakthrough infection followed by reinfection with BA.5/BF.7; BA.2 BTI+re: BA.2 breakthrough infection followed by reinfection with BA.5/BF.7; BA.5 BTI: BA.5 breakthrough infection, BF.7 BTI: BF7 breakthrough infection. D) The ACE2 binding affinity score of escape mutations located in 12 RBD epitopes. Possible RBD mutations encompassed those caused by single-step nucleotide changes on the BA.5.2 genome (EPI_ISL_16614729). The center line indicates the median, the box represents the interquartile range (IQR), the whiskers extend to the furthest data point in each wing that is within 1.5 times the IQR, and the dots represents outliers. Bonferroni adjusted p-values were calculated using the Wilcoxon rank sum test. ns: not significant.

